# Role of PARylation and PTEN Mutation on PARP and PARG Inhibitor Efficacy on Glioblastoma

**DOI:** 10.1101/2020.06.30.180216

**Authors:** Henre Hermanowski, Barbara Huebert, Christopher Aldrighetti, Jenna K Hurley, Delphine Quénet

## Abstract

Glioblastoma (GBM) is the most aggressive primary adult brain tumor, with a median survival of approximately 15 months. Despite novel therapeutic approaches, median survival has remained largely unchanged since the standard of care therapy for GBM was established nearly 15 years ago. Phosphatase and tensin homolog (PTEN) is a prognostic biomarker of GBM. PTEN mutation is associated with defects in homologous recombination (HR), making it a candidate for targeted therapy by synthetic lethality (SL). The SL concept has been clinically validated in HR-deficient breast and ovarian cancers upon treatment with poly(ADP-ribose) polymerase (PARP 1) inhibitors (PARPi). This inhibitor, as well as poly(ADP-ribose) glycohydrolase (PARG) inhibitors (PARGi), dysregulate PARylation post-translational modification, which plays a major role in DNA repair and genomic stability. To determine whether PARPi/PARGi promotes SL in GBM, this study investigated the effects of PARPi (veliparib and olaparib) and PARGi in GBM cells with wildtype versus mutant PTEN. Sensitivity to these drugs was analyzed in function of PTEN status. Specifically, PTEN-wildtype cells displayed higher levels of DNA damage after PARPi treatment compared to PTEN-mutant cells. However, focusing on DNA double-strand break (DSB) repair, there was no indication of efficient activation of non-homologous end joining (NHEJ) or homologous recombination (HR). These findings highlight the complex relationship between PARylation and PTEN. Thus, our results do not support the SL between PARP/PARG inhibition and PTEN mutations in GBM cells in absence of other DNA damaging agents.

## Introduction

Glioblastoma (GBM) is the most aggressive brain tumor, with a median survival of approximately 15 months, and accounts for more than half (55.1%) of all glioma subtypes [1–4]. Standard of care therapy for GBM, which has been established nearly 15 years ago, includes maximal surgical resection in conjunction with radiation therapy (60 Gy) and temozolomide (TMZ) chemotherapy (up to 6 cycles) [5]. However, this therapeutic strategy has yet to produce significant advances in median survival, largely due to the acquisition of TMZ resistance [6]. Despite the high demand for effective therapies, few options exist. Therapeutic efficacy is limited by the number of drugs that can cross the blood-brain barrier and remarkable intratumoral heterogeneity [7, 8].

The prognostic relevance of phosphatase and tensin homolog (PTEN) mutation or deletion, observed in 50-70% of GBM patients, remains debated. Several studies show that these genetic alterations may correlate with decreased survival, while others refute the role of PTEN alterations on survival [9–11]. These conflicting reports result from the insufficient characterization of individual PTEN mutations at the molecular level. This tumor suppressor has both phosphatase-dependent and phosphatase-independent activities that regulate myriad processes associated with metabolism, cellular proliferation, and genomic stability [12]. PTEN also modulates DNA double strand break (DSB) repair via the homologous recombination (HR) pathway, by debated mechanism(s) [12, 13]. The roles played by PTEN have opened the possibilities of a “BRCAness” phenotype that will induce synthetic lethality (SL) in PTEN-mutated GBM [14, 15].

PARylation is a transient, post-translational modification modulating numerous molecular mechanisms involved in the maintenance of genome stability, including DNA repair and transcription [16]. In the context of DNA damage, the polymer of poly(ADP-ribose) (PAR) is a cellular signal that participates in DNA damage accessibility, regulation of protein-protein interaction, and recruitment of integral DNA repair factors, acting as a chief modulator of genome integrity. In the presence of a DSB, poly(ADP-ribose) polymerase 1 (PARP1) promotes HR by competing with Ku70/Ku80 binding after break recognition, stimulating resection and RAD51 recruitment [17]. Interestingly, if resection is limited or unable to occur, alternative-non-homologous end joining NHEJ (alt-NHEJ), a minor and mutagenic DSB repair pathway dependent on PARP1, will be utilized. PAR degradation by poly(ADP-ribose) glycohydrolase (PARG) also contributes to the DNA damage response, facilitating the release of DNA repair machinery and helping to restore a functional chromatin structure [18]. The activities of PARP1 and PARG are involved in the recognition and signaling of DNA damage and the balance between different DNA repair pathways, highlighting the major role of PARylation in repair fidelity and genome stability [19]. Inhibitors of both PARP1 (PARPi) and PARG (PARGi) have been generated for their potential therapeutic interest [20, 21]. Most efforts have focused on PARPi, with the design of two types of inhibitors [21, 22]. Type 1 inhibitors (e.g., veliparib) compete directly with the NAD^+^ substrate for PARP1, inhibiting its enzymatic activity [21–23]. In contrast, type 2 inhibitors (e.g., olaparib) stabilize PARP-DNA complexes via a mechanism called “PARP1 molecular trapping”, which induces additional damage by collision with the transcription or replication machinery [21–23]. Type 2 inhibitors are significantly more cytotoxic than the genetic deletion of PARP1 itself, indicating that inactivated and trapped PARP1 produces direct DNA damage within the cell.

The potent PARGi PDD00017273 has recently been designed, yet its therapeutic efficacy has not yet been investigated [20]. *In vitro* studies show that this PARGi appears to sensitize HR-deficient cells (*e.g.*, BRCA1 mutated cells) via a different but unidentified molecular pathway than the PARPi olaparib [24, 25]. PARGi treatment leads to the accumulation of aberrant mitotic cells, likely a result of unrepaired DNA damage and stalled replication forks.

While PARP1 and PARG inhibition dysregulates the same post-translational modification (*i.e.,* PARylation), the downstream consequences of each are different, likely due to their specific roles in DNA repair [20]. Still, both PARPi and PARGi are postulated to sensitize DNA damage-response deficient cells following the concept of SL. To investigate the outcomes of inhibiting PARP1 and PARG in GBM, we compared the sensitivity and DNA damage response of two PARPi and one PARGi in GBM cells. We show that PTEN-mutated GBM cells are more sensitive to PARGi, and wildtype PTEN cells to PARPi. This difference of response is associated with PTEN and PARP1 expression levels. Despite the accumulation of cells in G2 in response to PARPi and PARGi, no cell death was observed in tested conditions. Interestingly, PARPi and PARGi induce the accumulation of DNA damage, in absence of other genotoxic agent, in PTEN-mutated GBM cells and wildtype PTEN cells, respectively. This work indicates that PTEN loss does not sensitize GBM cells to PARPi like BRCA does, but to PARGi, suggesting a functional link between dePARylation and PTEN.

## Methods

### Drugs

PARPi Veliparib (#S1004, Selleckem) and Olaparib (#S1060, Selleckem), and PARGi PDD00017273 (#5952, Tocris) were dissolved in 100% DMSO at 20mM and stored at −20°C.

### Cell Culture and Treatment

Three established GBM cell lines, LN-229 (PTEN-wildtype), U-87MG and U-118MG (PTEN mutated with the substitution mutation c.209+1G>T and c.1026+1G>T, respectively), were cultured in Dulbeco’s Modified Eagle’s Medium supplemented with 10% fetal bovine serum and 1% penicillin/streptomycin in a 5% CO_2_ tissue culture incubator at 37°C.

GBM cells were either untreated (NT, DMSO), or treated with the PARPi veliparib (IC_50_ concentration: 250 μM for LN-229 and 500 μM for U-87MG and U-118MG), PARPi olaparib (IC_50_ concentration: 10 μM for LN-229 and 100 μM for U-87MG and U-118MG), or PARGi PDD00017273 (IC_50_ concentration: 350 μM for LN-229 and 150 μM for U-87MG and U-118MG) for 24 hours. As a positive DNA damage control, cells were treated with 10 μM doxorubicin (#S1208, Selleckem) for 30 min in γH2AX and 53BP1 experiments and 4 hrs in Ku80 experiments. Cells were treated with 5 μM etoposide (#S1225, Selleckem) for 4 hrs in RAD51 experiments, and 2 mM hydroxyurea (#400046, Millipore) for 24 hrs in FANCD2 experiments. As an apoptotic control, cells were treated with 30 μg/mL digitonin (#300410, Millipore) for 30 min.

### Clonogenic Assay

Two hours after seeding on a P60 plates, 1,500 GBM cells were treated with DMSO, PARPi veliparib, PARPi olaparib or PARGi for 24 hrs. After 1X PBS wash, cells were grown for 2 weeks. Then, cells were fixed with 3.7% formaldehyde (#15680, EMS) and stained with 0.5% crystal violet (#C6158, Sigma) and colony formation assessed by manual counting. Three biological replicates with three technical replicates were performed.

### RNA extraction and analysis of gene expression via quantitative PCR

GBM cells were washed twice with cold 1X PBS, and RNAs were extracted with Trizol (#15596026, Ambion) following manufacturer’s instructions. RNAs were precipitated with isopropanol, DNase I treated and concentrated using the RNA Clean & Concentrator™ −5 Kit (#R1014, Zymo Research). Total RNA (1μg) was reverse transcribed using the SuperScript IV Kit (#1809150, Invitrogen) following manufacturer’s instructions. As a control, one reaction was performed in the absence of reverse transcriptase. qPCR was performed with 1X Sybr® Green Supermix (#172-5124, Bio-Rad) following manufacturer’s instructions. The plate was loaded with a QIAgility Robot (Qiagen) and run on 7500 Fast Real-Time PCR System (ThermoFisher) with the following standard conditions: (i) initial denaturation at 95°C for 2min, (ii) amplification over 40 cycles at 95°C for 15s, annealing at variable temperatures depending on primer set (see Table 1 for primer sequences) for 1min, and (iii) an elongation step at 95°C for 15s, 60°C for 1min, 95°C for 15s, and 60°C for 15s. Levels of RNA expression were determined against a standard curve, and ratios [gene of interest/rRNA 18S] were assessed (rRNA 18S primers kindly provided by Dr. B. Girard, University of Vermont Larner College of Medicine). Four biological replicates of cDNA and two technical replicates of the standard curve were run for each gene of interest. Two-sided t-tests were used to determine the significance of expression between cell lines. Information on the plasmids to generate the standard curve are available upon request.

**Table 1:**
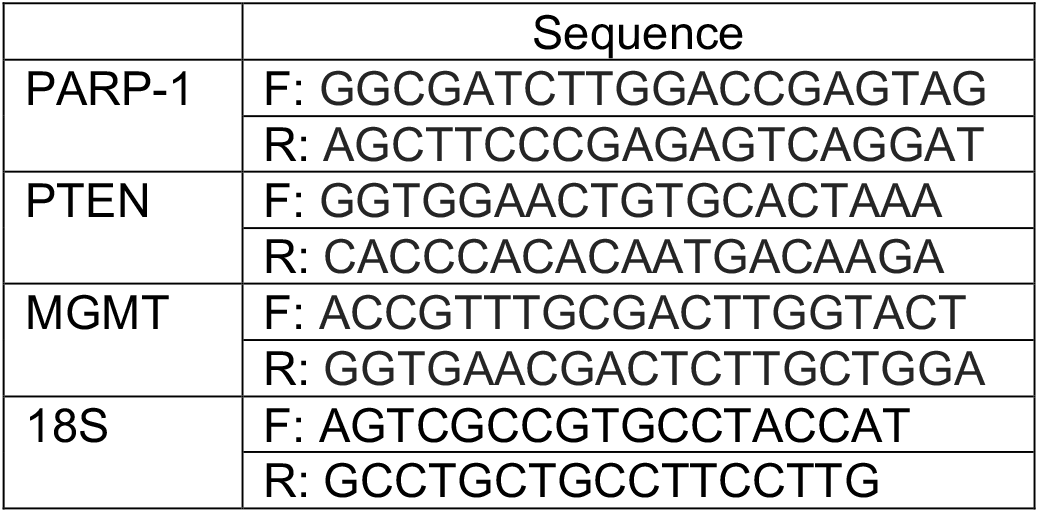
List of primer sets.

### Western Blotting

GBM cell lines were washed twice with cold 1X PBS before being collected in RIPA buffer (50 mM Tris pH 7.4, 150 mM NaCl, 0.1% SDS, 0.5% sodium deoxycholate(#102906, MP Biomedicals), 1% Nonidet™ P-40 substitute (#74385, Sigma-Aldrich), 1X cocktail protease inhibitor (#11873580001, Roche)). After quantification by Bradford assay (#500-0205, Bio-Rad), proteins were separated on a 10% SDS-PAGE, and then, transferred onto a nitrocellulose membrane (#1620112, Bio-Rad). The membrane was blocked with 1X PBS, 1% BSA (#A3294, Sigma-Aldrich) for 60 minutes at room temperature and incubated overnight at 4°C with appropriate primary antibody (Table 2) diluted in 1X PBS, 0.5% BSA, 0.1% Tween 20 (#BP337-550, Fisher Scientific). After 3 washes in 1X PBS, 0.01% Tween 20 for 5 min at room temperature, the membrane was incubated for 60 minutes at room temperature with the secondary antibody (Table 2) diluted in 1X PBS, 0.5% BSA, 0.1% Tween 20. Membranes were washed 3 times with 1X PBS, 0.01% Tween 20 and digitally scanned using a LiCor Odyssey® Infrared Scanner. Band signal intensities were quantified using the open source program ImageJ (https://imagej.nih.gov/ij/). Signal ratios of PAR or PARG to β-actin were determined. All experiments were performed at least in triplicate. Statistical significance was assessed by student’s t-test using mean and standard deviation. P-values below 0.05 were considered significant.

**Table 2:**
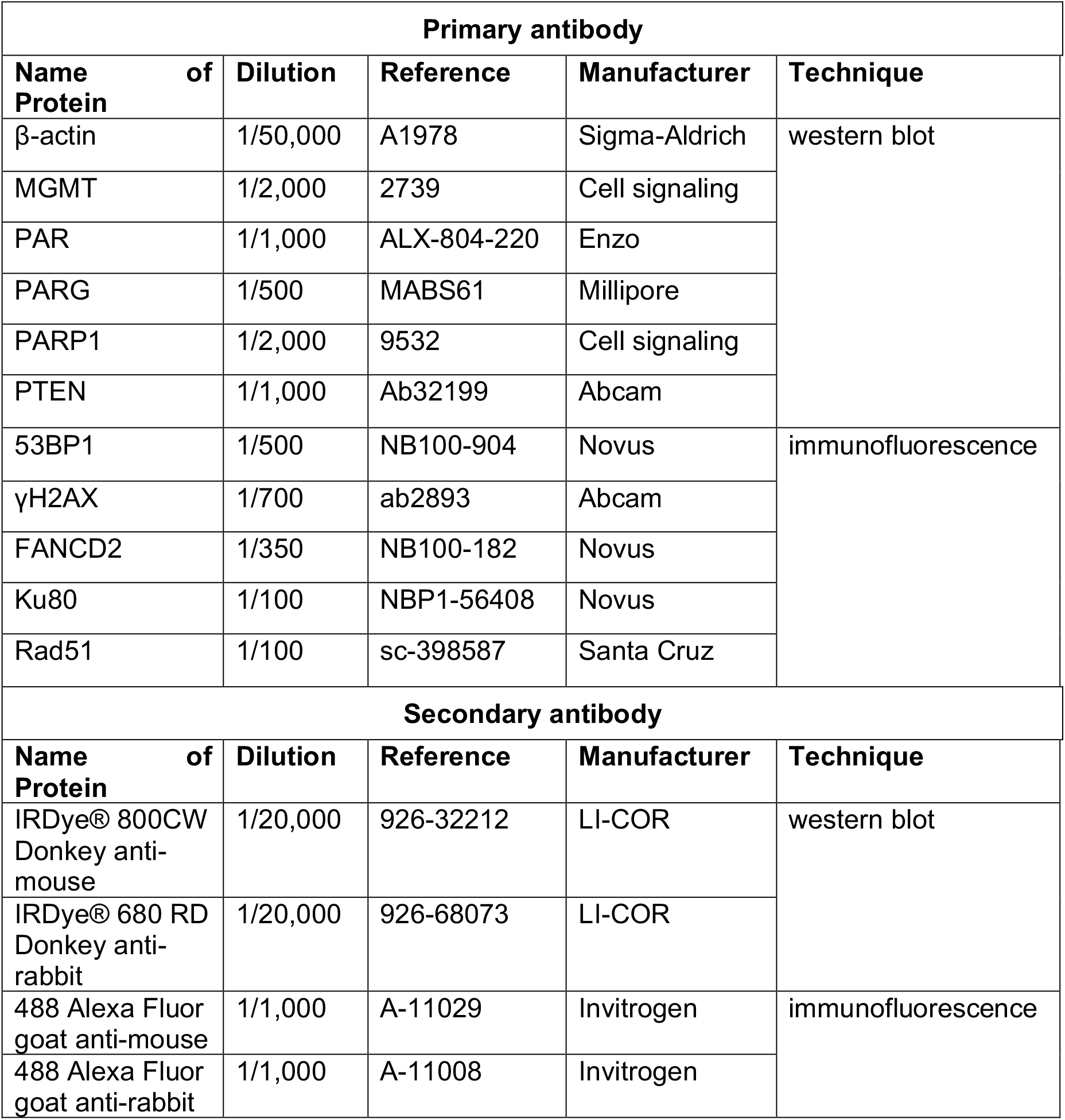
List of primary and secondary antibodies.

### Flow Cytometry Analysis

To study cell cycle distribution, GBM cells were treated with TMZ, PARPi veliparib, PARPi olaparib, or PARGi as described in *Cell Culture and Treatment*. In addition, cells were treated with digitonin to induce apoptosis or doxorubicin to generate DNA damage. Then, cells were collected, PBS-washed, fixed with 1% paraformaldehyde (PFA, #50-980-494, EMS) and permeabilized in ice-cold 70% ethanol. Cell cycle distribution of cells resuspended in propidium iodide solution (50 μg/mL PI [#53-705-9100MG, Millipore]; 0.1% Triton X-100; 1X PBS) was analyzed by flow cytometry on the BD LSRII equipment available at the Harry Hood Bassett Flow Cytometry and Cell Sorting Facility. Based on cellular DNA content, cells were categorized as Sub-G1, G1, S, G2 and polyploid.

To study cell death, cells were treated with digitonin, PARPi veliparib, PARPi olaparib, or PARGi as described in *Cell Culture and Treatment*. Apoptosis and necrosis were analyzed using the PI/Annexin V staining kit (#V13241, ThermoFisher), following manufacturer’s instructions. Cells were analyzed by flow cytometry. Cell populations were categorized into one of three groups: living cells (PI^−^/annexin V^−^), necrotic and apoptotic cells (PI^+^/annexin V^+^), and early apoptotic cells (PI^−^/annexin V^+^).

Two and three biological replicates were performed for the cell cycle and cell death assays, respectively, with 10,000 cells counted per condition. Data were acquired and analyzed using FlowJo software. Standard deviations were determined and two-way ANOVA tests were performed to assess the significance of the results using Prism 8 software (Graph Pad). P-values below 0.05 were considered significant.

### γH2AX, 53BP1, and FANCD2 Immunofluorescence

To study γH2AX, 53BP1 and FANCD2 expression, cells were treated with drugs as described in *Cell Culture and Treatment* prior to fixation in ice-cold 4% PFA, 2% sucrose, 1X PBS for 10 min. After two washes with 1X PBS for 5 min at room temperature, cells were permeabilized in 0.5% Triton X-100, 20 mM HEPES pH7.4, 50 mM NaCl, 3 mM MgCl_2_, 300 mM sucrose, 1X PBS for 5 min at room temperature. Cells were washed two times in 1X PBS, 0.1% Triton X-100 for 5 min at room temperature, and then incubated with primary antibody (Table 2) diluted in 1X PBS, 2% BSA at 4°C overnight in a humidified chamber. After two wash in 1X PBS, 0.1% Triton X-100, cells were incubated with secondary antibody (Table 2) diluted in 1X PBS, 2% BSA for 1 hr at room temperature in a dark humidified chamber. Finally, cells were washed three times with 1X PBS, 0.1% Triton X-100, stained with 4’,6-diamidino-2-phenylindole (DAPI, 0.1 μg/mL; #D1306, Invitrogen) and mounted with Prolong Gold antifade medium (#P10144, Invitrogen).

### Ku80 Immunofluorescence

To assess Ku80 expression, cells were treated with drugs as described in *Cell Culture and Treatment*. Cells were washed once with ice-cold 1X PBS, incubated twice in CSK buffer (10 mM PIPES pH6.8, 100 mM NaCl, 300 mM sucrose, 3 mM MgCl_2_, 1 mM EGTA) with 0.7% Triton X-100 and 0.3 mg/mL RNase A for 3 min, and then rinsed with 1X PBS at room temperature. Cells were fixed with 2% PFA in 1X PBS for 15 min, rinsed in 1X PBS, blocked with 1X PBS, 0.1% Tween 20, 5% BSA for 1 hr at room temperature, and incubated with the primary antibody (Table 2) diluted in 1X PBS, 0.1% Tween 20, 5% BSA at 4°C overnight in a humidified chamber. After three wash with 1X PBS, 0.1% Tween 20 for 5 min each, cells were incubated with secondary antibody (Table 2) diluted in 1X PBS, 0.1% Tween 20, 5% BSA for 1 hr in the dark at room temperature. Cells were washed in the dark at room temperature two times with 1X PBS, 0.1% Tween 20 for 5 min, and once with 1X PBS containing DAPI (0.1 μg/mL) for 10 min. After two washes with 1X PBS for 5 min, cells were rinsed in distilled water and mounted on slides with Prolong Gold antifade medium.

### RAD51 Immunofluorescence

To study Rad51 expression, cells were treated with drugs as described in *Cell Culture and Treatment* prior to fixation with ice-cold 100% methanol for 20 min at 4°C. Cells were permeabilized with ice-cold 1X PBS, 0.5% Triton, then washed with ice-cold 1X PBS for 5 min on ice, blocked with 1X PBS, 0.1% BSA for 1 hr on ice, and incubated with primary antibody (Table 2) diluted in 1X PBS, 0.1% Tween 20, 5% BSA at 4°C overnight in a humidified chamber. Cells were washed three times for 5 min with 1X PBS and incubated with secondary antibody (Table 2) diluted in 1X PBS, 0.1% Tween 20, 5% BSA for 1 hr in a dark humidified chamber at room temperature. Cells were washed at room temperature in the dark with 1X PBS for 5 min and once with 1X PBS containing DAPI (0.1 μg/mL) for 10 min. Cells were washed with 1X PBS for 10 min in the dark at room temperature, rinsed in distilled water, and mounted on slides with Prolong Gold antifade medium.

### Immunofluorescence Analysis

Slides were observed under a Nikon Ti-E inverted microscope using NIS-Elements microscope imaging software. Three biological replicates were performed for each marker, with at least 100 cells counted per condition. Pictures were analyzed using the open source image processing program Fiji (https://imagej.net/).

For γH2AX and 53BP1, cells were manually counted and categorized as followed: ≤1 focus, 2-5 foci and ≥6 foci. For FANCD2, cells were either classified based on the absence or presence of foci. The percentages of cells within each category were represented on a 100% stacked column graph.

Ku80 signal intensity in the nucleus was measured automatically by the NIS-Elements microscope imaging software. The DAPI signal was used to identify the region of interest, then Alexa^488^ fluorescent signal intensity within that region was quantitated. For each biological replicate, intensity values were normalized to the untreated.

Nuclear Rad51 foci number was measured automatically by the NIS-Elements microscope imaging software as described above for Ku80. For each biological replicate, values were tabulated and graphed on a box plot chart where the average, median, minimum, maximum, and first and third quartiles are indicated.

Two-way ANOVA was performed to assess statistical variance between cell lines and/or treatment using Prism 8 software (Graph Pad). P-values below 0.05 were considered significant.

## Results

### PARP1 expression is decreased in PTEN-mutated GBM cells

To study a potential functional interaction between PARP1 and PTEN in GBM cells, mRNA and protein levels of these two factors, as well as the DNA repair enzyme O-6-methylguanine-DNA methyltransferase (MGMT), whose expression confers chemotherapy resistance, were assessed by qPCR and western blot, respectively. PTEN and PARP1 mRNA levels were higher in GBM cells expressing wildtype PTEN (LN-229) compared to PTEN mutated cells (Figure 1A). Expression of MGMT transcripts was almost undetectable in all GBM cells, indicating these cells would be responsive to chemotherapy. Then, we compared the expression profile of these three factors at the protein level (Figure 1B). Western blot analysis revealed similar expression patterns; PARP1 and PTEN were expressed in LN-229 cells, but barely detectable in U-87MG and U-118MG cells. These data suggest that PTEN and PARP1 expression may be co-regulated

**Figure 1:**
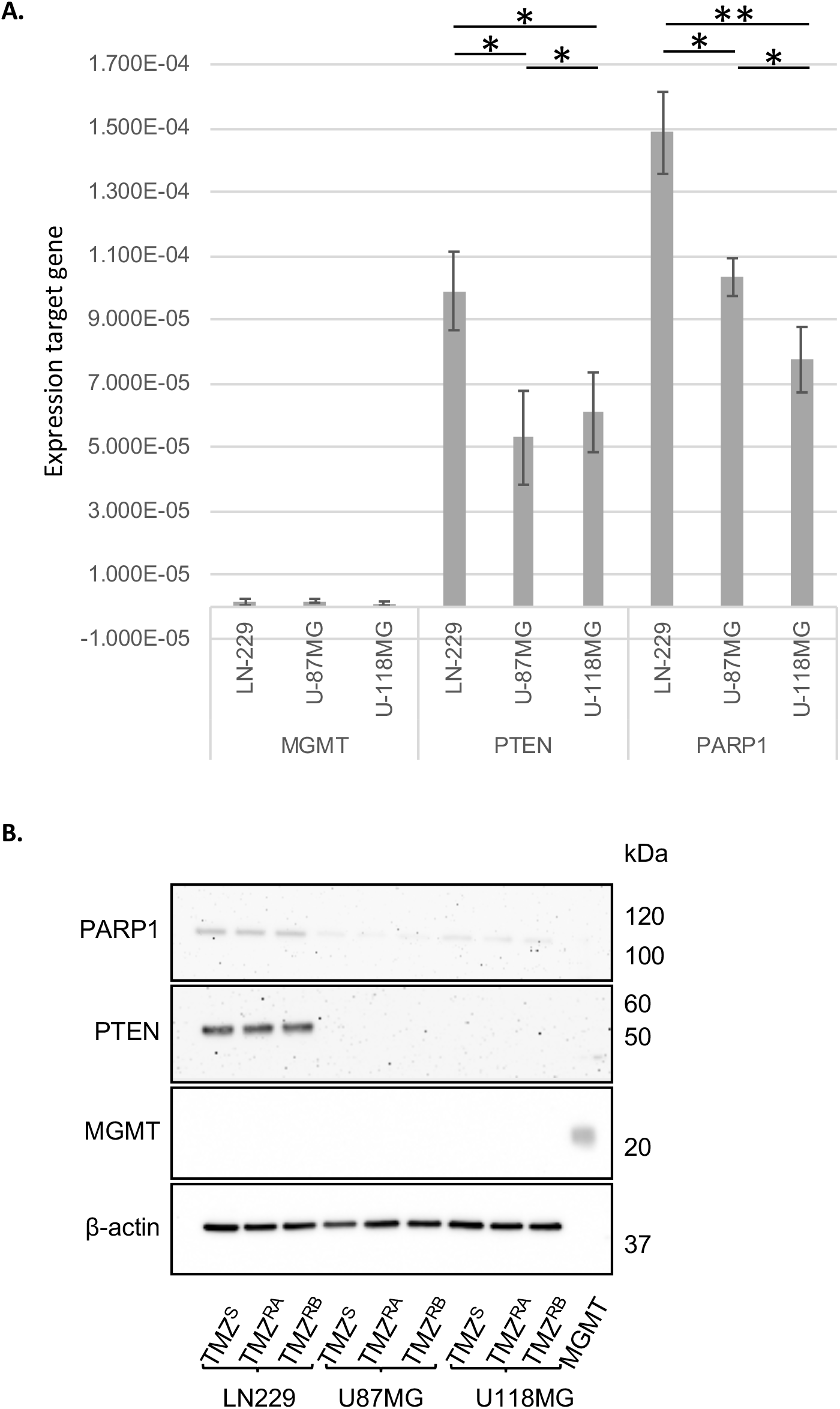
PTEN and PARP1 are co-expressed in LN-229 cells. A. Total RNAs were purified and expression level of gene of interest were analyzed by qPCR. Transcript level was determined by a standard curve and normalized to 18S rRNA. B. Total proteins were extracted from GBM cells and analyzed by western blot.

### PTEN-wildtype and PTEN-mutant GBM cells are more sensitive to PARPi and PARGi, respectively

To analyze their sensitivity to PARPi and PARGi, GBM cells were incubated for 24 hrs with the PARPi veliparib, PARPi olaparib or a PARGi. Colony formation of treated cells relative to vehicle not-treated controls was assessed after two weeks (Figure 2). This assay allowed the determination of an approximative IC_50_ for each tested drug: PARPi veliparib (IC_50_ concentration: 250 μM for LN-229 and 500 μM for U-87MG and U-118MG), PARPi olaparib (IC_50_ concentration: 10 μM for LN-229 and 100 μM for U-87MG and U-118MG), PARGi PDD00017273 (IC_50_ concentration: 350 μM for LN-229 and 150 μM for U-87MG and U-118MG). These data showing LN-229 cells, which express wildtype PTEN, are more sensitive to PARPi, while U-87MG and U-118MG cells, which express mutant PTEN, are more sensitive to PARGi further support a potential functional relationship between PTEN and PARP1.

**Figure 2:**
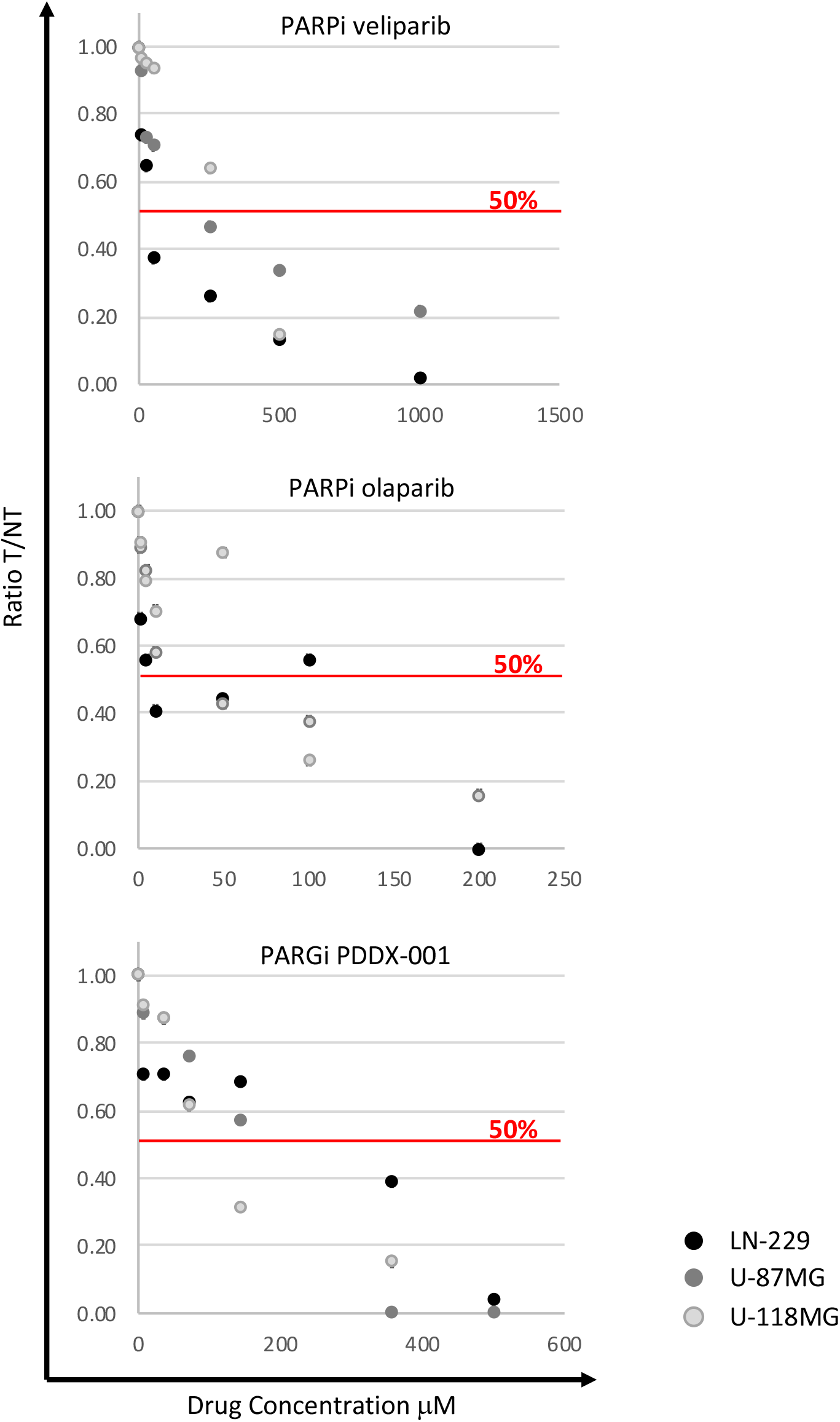
Sensitivity to PARPi and PARGi is dependent on the expression of PTEN. GBM cells were treated with increasing concentrations of PARPi (veliparib or olaparib) or PARGi PDD00017273 for 24 hours. Surviving clones were counted after staining with crystal violet to determine the IC_50_. Graphs represents the colony ratio [Treated/not-treated] as a function of drug concentration.

### PARPi in LN-229 cells and PARGi in U-87MG and U-118MG cells preferentially leads to G2 arrest

To analyze the overall cellular effect of PARP1 and PARG inhibition (24 hrs treatment at IC_50_) in GBM cells, the cell cycle progression of unsynchronized LN-229, U-87MG and U-118MG cells was studied by flow cytometry (Figure 3). Non-treated LN-229 (NT) cells were mainly in the G1 phase (~56%), while ~23% and ~15% of cells were in S and G2 phases, respectively (Figure 3). Few LN-229 cells were in sub-G1 or displayed polyploidy (<5% each). This same pattern was observed upon treatment with digitonin (apoptotic-inducing agent), doxorubicin (DNA-damaging agent) and TMZ (DNA-damaging GBM chemotherapeutic agent). In contrast, LN-229 cells treated with the PARPi veliparib or olaparib were arrested in the G2 phase (P<0.0001 compared to NT), while PARG inhibition led to the accumulation of LN-229 cells in the G1 phase (P=0.0012 compared to NT).

**Figure 3:**
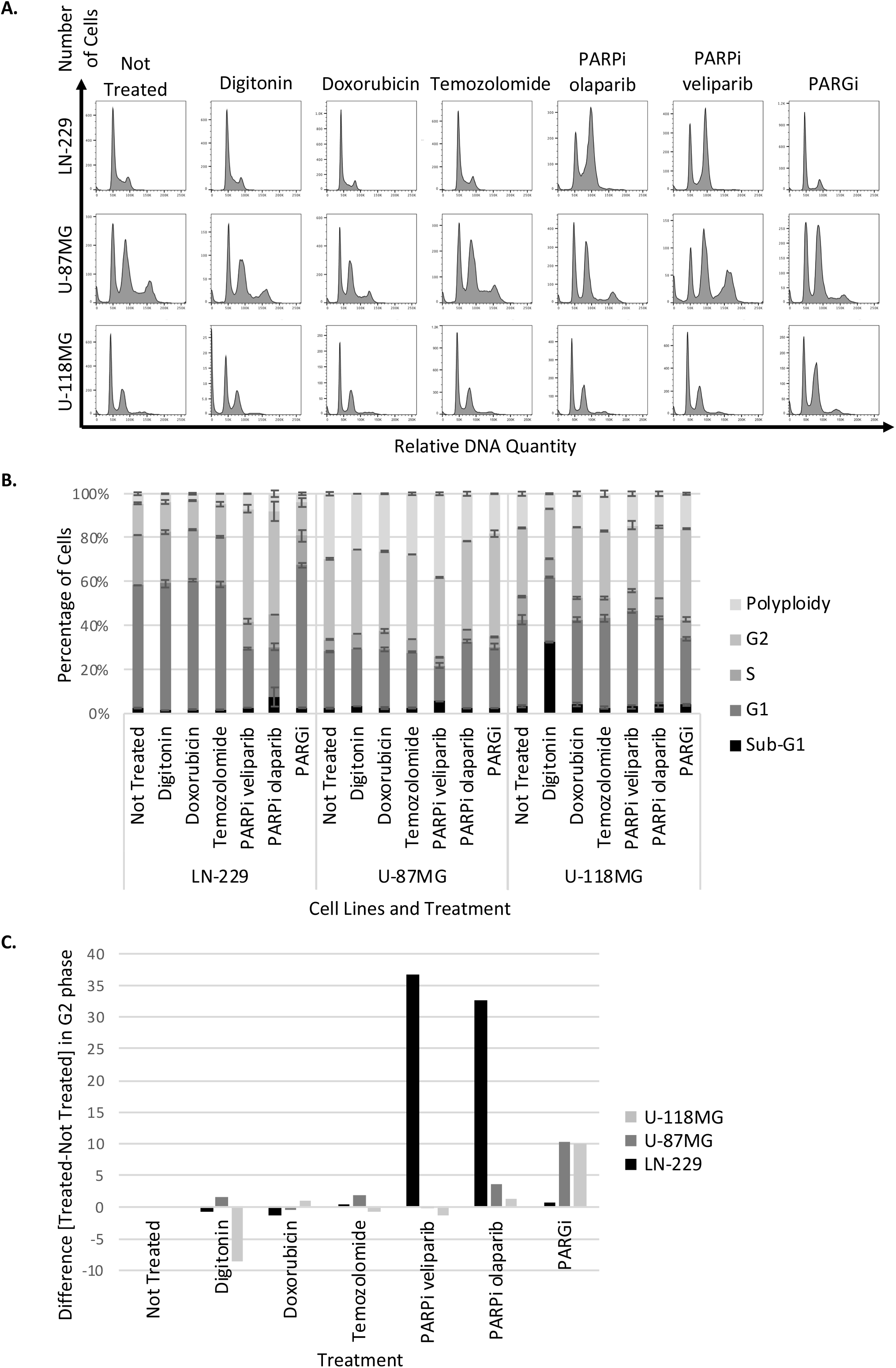
PARPi and PARGi induce G2 arrest in GBM cells. GBM cells were treated with PARPi (veliparib or olaparib) or PARGi PDD00017273 at their IC_50_ for 24 hours. Cells were fixed and stained with propidium iodide before cell cycle analysis by flow cytometry. DNA quantity was measured to place the cells into 5 categories (Sub-G1, G1, S, G2, and Polyploidy). A) Cell cycle distribution as a function of the relative DNA quantity was plotted for one representative experiment. B) The average percentage of cells observed in each phase of the cell cycle was plotted as a function of the cell line and treatment. Error bars correspond to standard deviation. C) The difference in percentage between treated and non-treated cell populations in the G2 phase was represented for each treatment. Digitonin, doxorubicin, and temozolomide were used as positive controls. Data are representative of two biological replicates.

U-87MG cells expressing mutant PTEN showed different growth arrest patterns that were PARPi specific. U-87MG NT cells were preferentially in G1 and G2 phases (~26% and ~37%, respectively) compared to S phase and sub-G1 (<6% each) (Figure 3). Moreover, U-87MG NT cells displayed high levels of polyploidy (~30%). Digitonin, doxorubicin and TMZ treatments did not significantly change this pattern. Intriguingly, the PARPi veliparib was associated with increased polyploidy (P<0.0001 compared to NT) and decreased G1 cell population (P<0.0001 compared to NT), while the PARPi olaparib resulted in decreased polyploidy (P<0.0001 compared to NT) and increased G1 cell population (P<0.0001 compared to NT). There was a significant increase in the G2-U87-MG cell population after PARGi treatment (Figure 3C).

The cell cycle profile of U118-MG NT cells was similar to U-87MG NT cells: ~3% in sub-G1, ~40% in G1, ~10% in S, ~31% in G2 and ~16% polyploidy (Figure 3). The same pattern was observed after doxorubicin, TMZ and PARPi treatment. In contrast, treatment with digitonin caused a significant increase in the sub-G1 population as compared to NT cells (P<0.0001). Veiiparib and olaparib treatments did not affected the cellular distribution of U118-MG cells. In contrast to PARPi, PARGi was associated with an accumulation of cells in G2 (Figure 3C; P<0.0001 compared to NT). The preferential G2 phase accumulation of LN-229 PTEN-wildtype cells upon PARP inhibition and U-87MG and U-118MG PTEN-mutant cells after PARGi treatment suggests a common mechanism of PARPi and PARGi independent of PTEN expression.

### PARPi and PARGi treatment does not induce GBM cell death

To determine whether PARPi- and/or PARGi-associated G2 arrest triggered cell death in GBM cells based on PTEN status, markers of apoptosis and necrosis were analyzed by flow cytometry (Figure 4). As a positive control, GBM cells were treated with digitonin, an apoptotic/necrotic-inducing agent. Cells were positive for both annexin V and PI, indicating that they were dead. In contrast, GBM cells treated with the DNA-damaging agent doxorubicin were negative for both markers, suggesting that the treatment was not toxic enough to induce cell death. Although PARPi and PARGi treated cells showed G2 growth arrest, neither PI nor FITC-annexin V signals were detected upon PARP or PARG inhibition, suggesting that 24 hr treatment at IC_50_ was not sufficient to induce cell death by apoptosis or necrosis.

**Figure 4:**
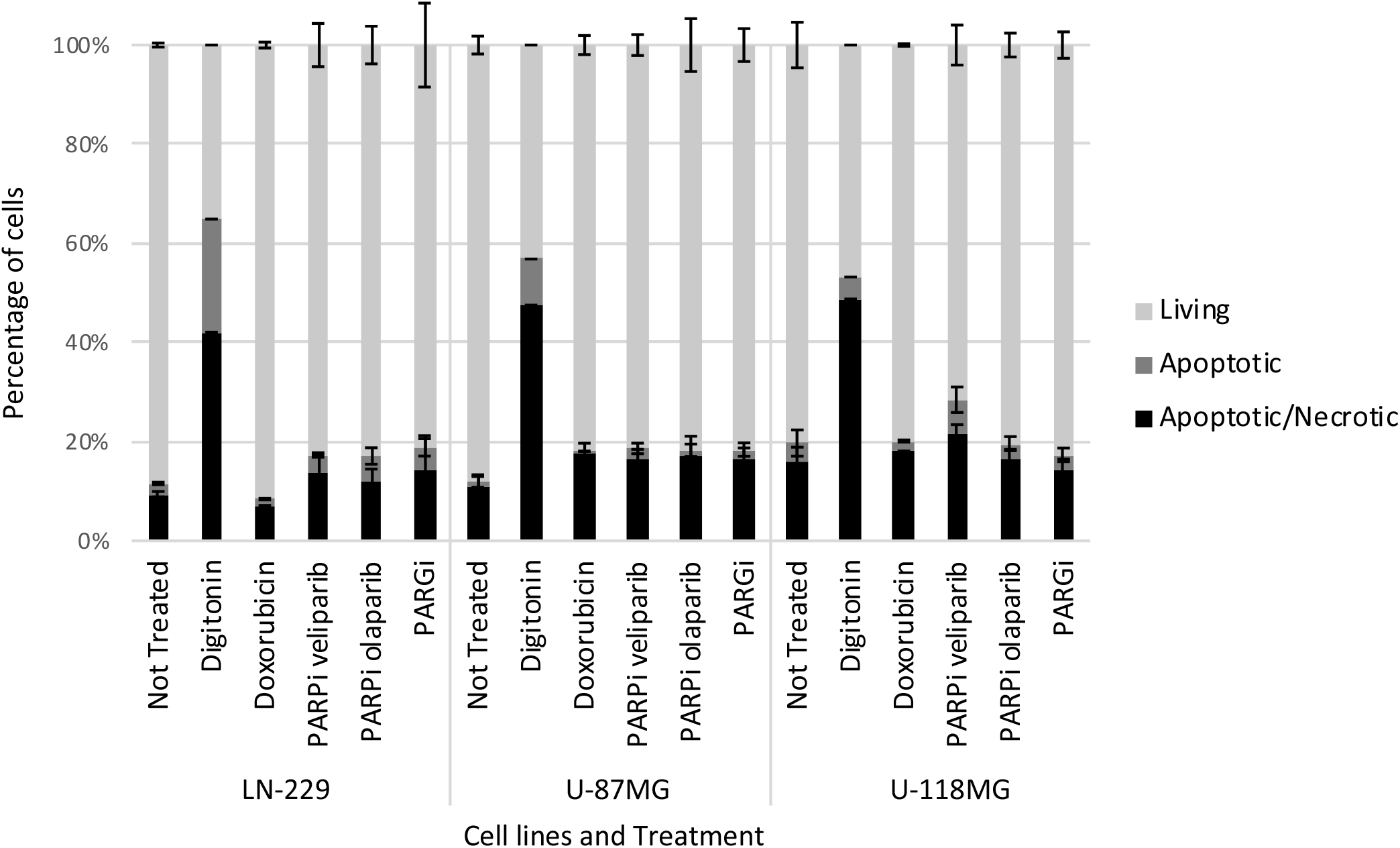
PARPi and PARGi do not induce apoptosis or necrosis in GBM cells. GBM cells were treated with PARPi (veliparib or olaparib) or PARGi PDD00017273 at their IC_50_ for 24 hours. Following annexin V and propidium iodide (PI) staining, apoptosis and necrosis activation was analyzed by flow cytometry. The average percentage of cells observed as apoptotic/necrotic (annexin V^+^, PI^+^; black), apoptotic (annexin V^+^; medium gray), or living (annexin V^−^, PI^−^; light gray) was plotted for each cell line and treatment. Digitonin was used as positive control. Error bars represent standard deviation. Data are representative of two biological replicates.

### PARPi triggered the accumulation of γH2AX foci in PTEN-wildtype GBM cells, while PTEN-mutant cells were less prone to DNA damage upon PARPi and PARGi

The accumulation of cells in the G2 phase can result from activation of the G2/M DNA damage checkpoint [26]. Arrest at the G2/M checkpoint permits examination of cells for the presence of DNA damage or incomplete DNA replication, thereby preventing compromised cells from entering mitosis [27]. To determine whether PARP1 or PARG inhibition induces a DNA damage response, the level of γH2AX foci, marker of DNA damage, was assessed.

The majority (~60%) of LN-229 NT cells displayed one or fewer γH2AX foci, while ~10% and ~25% exhibited 2-5 foci and ≥6 foci, respectively (Figure 5). As expected, doxorubicin treatment was associated with a significant increase in the presence of γH2AX foci (~80% of cells with ≥6 foci, P<0.0001 compared to NT). A similar increase in γH2AX foci was observed after PARP1 inhibition with veliparib and olaparib (P<0.0001 compared to NT). In contrast, PARG inhibition did not significantly increase the presence of γH2AX foci. This suggests that the DNA damage generated by PARPi in U87-MG GBM cells may be PTEN-dependent.

**Figure 5:**
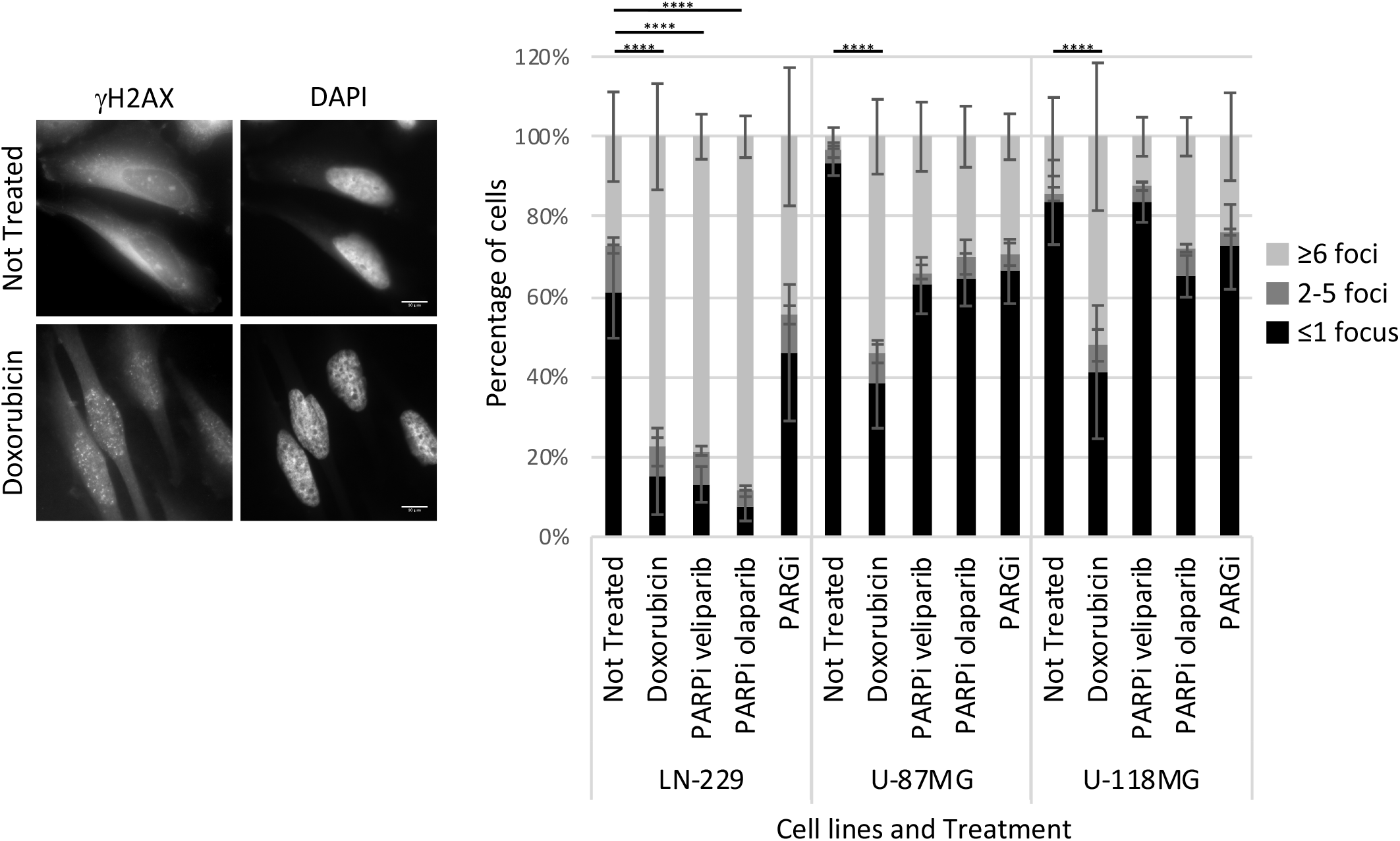
PARPi and PARGi generate DNA damage in PTEN-wild type GBM cells. After 24 hours of treatment with either PARPi (veliparib or olaparib) or PARGi PDD00017273 at their IC_50_, GBM cells were fixed and immunofluorescence-stained with γH2AX. Nuclei were stained with DAPI. As a positive control, cells were treated with doxorubicin. Representative images are presented (left panel). At least 100 cells displaying either ≤1 γH2AX focus (black), 2-5 γH2AX foci (medium gray) or ≥6 γH2AX foci (light gray) were counted per condition, per experiment. The average percentage of cells in each category is plotted for each cell line and treatment. Error bars represent standard deviation. Statistical significance was assessed by ANOVA. Data are representative of three biological replicates. **** P<0.0001.

In contrast to LN-229 cells, a lower percentage of U-87MG NT cells displayed γH2AX foci (~5% of cells with ≥6 foci, Figure 5). However, γH2AX-positive cells increased after doxorubicin treatment to ~60% (P<0.0001 compared to NT). PARPi and PARGi did not significantly increase the number of γH2AX foci compared to NT cells (~40% for PARPi veliparib, ~30% for PARPi olaparib and ~30% for PARGi for cells with ≥6 foci).

Similar to U-87MG cells, U-118MG NT cells presented few γH2AX foci (~10% of cells with ≥6 foci, Figure 5), but doxorubicin treatment significantly increased the presence of γH2AX foci (~50% of cells with ≥6 foci, P<0.0001 compared to NT). Similarly, PARP or PARG inhibition did not significantly increase the number of γH2AX-positive cells (PARPi veliparib: ~15%, PARPi olaparib: ~25%, PARGi: ~20%). These data suggest that PTEN-wildtype GBM cells, but not PTEN-mutant GBM cells, accumulate DNA damage upon PARP inhibition, further suggesting a link between PTEN and PARP1.

### PARP inhibition was associated with 53BP1 foci formation in PTEN-wildtype but not PTEN-mutant GBM cells

The ability of GBM cells to utilize HR versus NHEJ was analyzed by measuring the frequency of 53BP1 foci formed upon PARPi or PARGi treatment using an immunofluorescent approach (Figure 6A).

**Figure 6:**
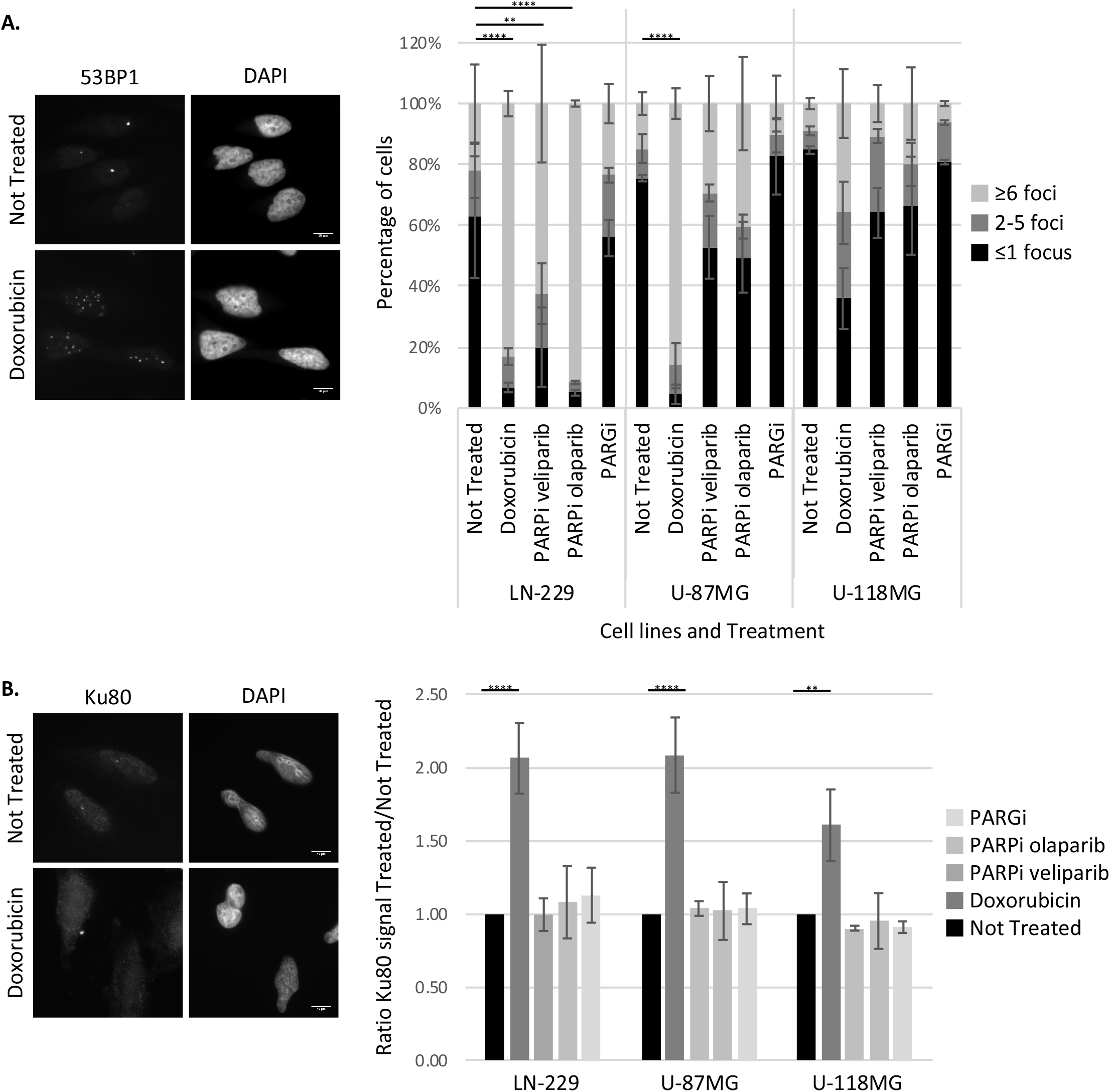
PARPi and PARGi activate NHEJ in PTEN-wildtype GBM cells. After 24 hours of treatment with either PARPi (veliparib or olaparib) or PARGi PDD00017273 at their IC_50_, GBM cells were fixed and immunofluorescence-stained for 53BP1 or Ku80. Nuclei were stained with DAPI. As a positive control, cells were treated with doxorubicin. Representative images are presented (left panel). A. At least 100 cells displaying either ≤1 53BP1 focus (black), 2-5 53BP1 foci (medium gray) or ≥6 53BP1 foci (light gray) were counted per condition, per experiment. The average percentage of cells in each category was plotted for each cell line and treatment. Error bars represent standard deviation. Statistical significance was assessed by ANOVA. Data are representative of three biological replicates. **** P<0.0001; ** P≥0.0021. B. Nuclear signal intensity of Ku80 was assessed by analyzing at least 100 cells. The ratio of Ku80 signal in the treatment group to the Not Treated group was plotted for each cell line and treatment. For both panels, error bars represent standard deviation. Statistical significance was assessed by ANOVA testing. Data are representative of three biological replicates. **** P<0.0001; ** P≥0.0021.

Only ~20% of LN-229 NT cells displayed ≥6 53BP1 foci. However, doxorubicin treatment increased this percentage to ~80% (P<0.0001 compared to NT). Interestingly, more 53BP1-positive cells were present following PARP inhibition with olaparib than veliparib (~90% *vs.* ~60% of cells with ≥6 foci, P<0.0001 for olaparib and P=0.0066 for veliparib compared to NT). Similar to no increase in γH2AX foci in PARGi treated LN-229 cells (Figure 5), PARGi did not induce the formation of 53BP1 foci.

In U-87MG NT cells, 53BP1 foci were observed in ~15% of cells, and ~90% of cells in response to doxorubicin treatment (P<0.0001 compared to NT) (Figure 6A). PARP1 inhibition did not significantly increase the number of 53BP1 foci (~25% for veliparib and ~40% for olaparib for cells with ≥6 foci). PARGi did not induce the formation of 53BP1 foci in U-87MG cells (~10% of cells with more than 6 foci), similar to that observed for LN-229 cells.

While only ~10% of U-118MG NT cells displayed ≥6 53BP1 foci compared to ~40% of U-118MG doxorubicin-treated cells, this difference was not statistically significant (Figure 6A). Similarly, neither PARPi nor PARGi treatment increased the percentage of 53BP1 foci (~10% for veliparib, ~20% for olaparib and ~5% for PARGi of cells with ≥6 foci).

These data indicate that upon PARPi treatment, the NHEJ marker 53BP1 forms foci in PTEN-wildtype GBM cells, but not in PTEN-mutant GBM cells. Not surprisingly, PARGi did not lead to 53BP1 foci formation either in wildtype or mutant PTEN GBM cells, potentially due to the absence of DNA damage (Figure 5).

### PARP and PARG inhibition did not significantly alter nuclear localization of Ku80 in GBM cells

To examine the potential activation of NHEJ repair pathway in PTEN-wildtype GBM cells after PARP1 inhibition, the nuclear localization of Ku80, a NHEJ mediator, was quantified immunocytochemically (Figure 6B). In contrast to γH2AX and 53BP1, Ku80 does not form foci, but its expression level significantly increases in the nucleus upon DNA damage. Thus, nuclear Ku80 signals were quantified following treatment and compared to the NT condition (ratio: nuclear Ku80 signal “Treated” / nuclear Ku80 signal “Not Treated”). As expected, the Ku80 ratio increased upon doxorubicin treatment in all three GBM cell lines, indicating that DNA damage can be repaired by a Ku80-dependent pathway in these cells (P<0.0001 compared to NT). However, PARPi and PARGi treatment did not significantly increase the nuclear signal of Ku80, suggesting that NHEJ may not be the prime pathway to repair PARPi- or PARGi-induced damage.

### RAD51 foci accumulate in PTEN-wildtype GBM cells after PARP inhibition, and in PTEN-mutant GBM cells after PARPi olaparib and PARGi treatment

To measure HR functionality in response to PARPi and PARGi in GBM cells RAD51 foci formation was analyzed (Figure 7, supplemental table 2). The number of RAD51 foci ranged from 0 to >120 in function of the treatment.

**Figure 7:**
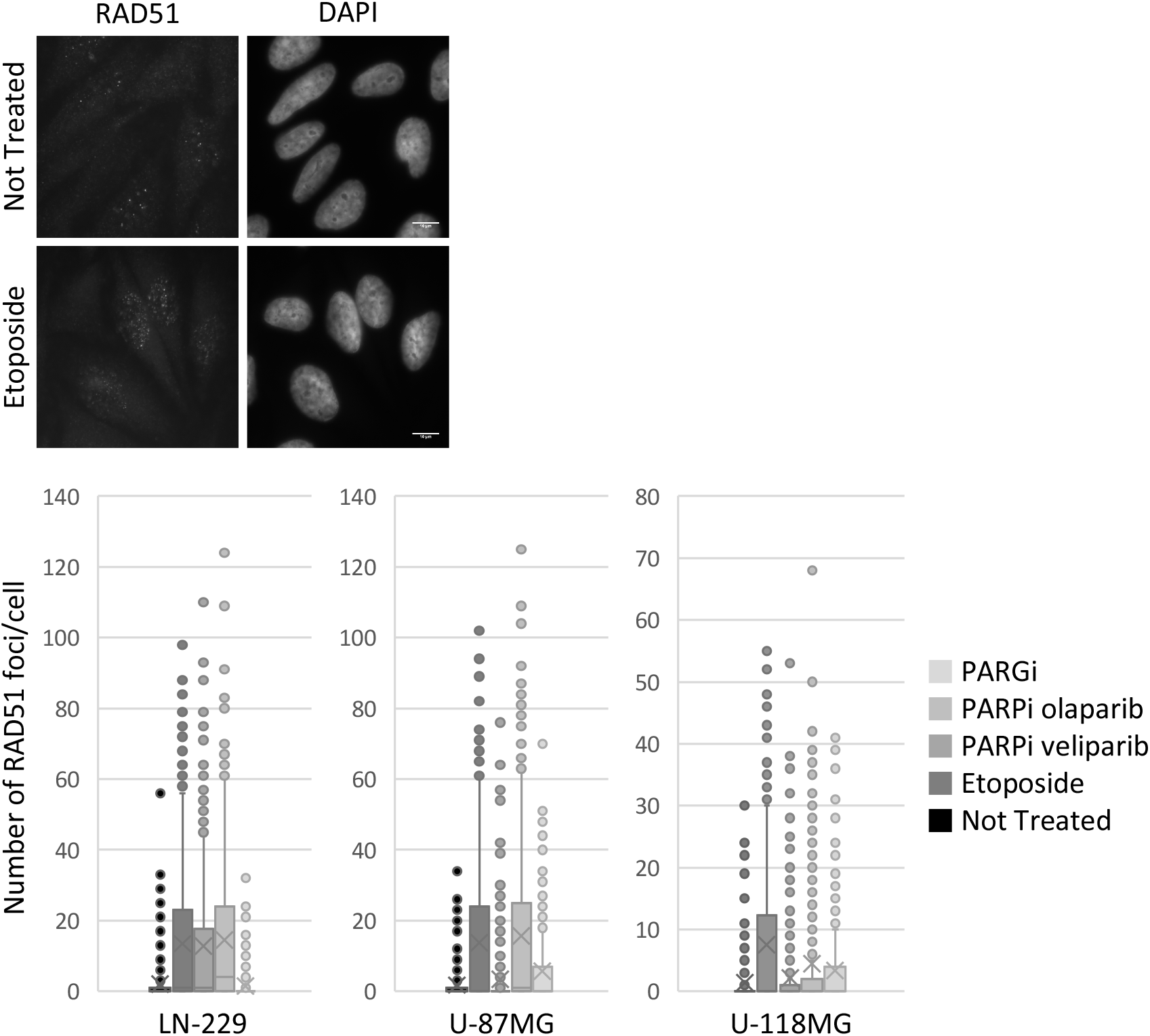
PARPi and PARGi activate HR in both PTEN-wild type and PTEN-mutant GBM cells. After 24 hours of treatment with either PARPi (veliparib or olaparib) or PARGi PDD00017273 at their IC_50_, GBM cells were fixed and immunofluorescence-stained for RAD51. Nuclei were stained with DAPI. As a positive control, cells were treated with etoposide. Representative images are presented (left panel). RAD51 foci number was assessed by counting at least 100 cells per condition, per experiment. The number of foci per cell was plotted for each cell line and treatment. The box plot indicates the average, median (x), minimum, maximum, and first and third quartiles. Outliers are depicted by dots beyond the interquartile range. Statistical significance was assessed by ANOVA. Data are representative of three biological replicates.

LN-229 NT cells displayed few RAD51 foci relative to cells treated with etoposide (mean ~2 *vs.* ~13 RAD51 foci, respectively, P<0.0001, Figure 7). Both PARPi veliparib and olaparib treatments led to increased RAD51 foci (P<0.0001 compared to NT). In contrast, PARGi-treated cells displayed a similar number of RAD51 foci compared to non-treated LN-229 cells. These data suggest that PARPi, but not PARGi, activates the HR pathway in PTEN-wildtype GBM cells.

As in LN-229 cells, few RAD51 foci were present in U-87MG NT cells (mean ~2, Figure 7). As expected, this number increased after etoposide treatment (mean ~14 RAD51 foci, P<0.0001 compared to NT). While PARPi veliparib was not associated with an overall increase of RAD51 foci, PARPi olaparib did significantly increase RAD51 foci (mean ~3 *vs.* ~16 RAD51 foci, respectively, P<0.0001 compared to NT). PARGi also increased the mean RAD51 foci, but to a lesser extent (mean ~6, P=0.0078 compared to NT).

U-118MG NT cells did not display detectable RAD51 foci, but etoposide treatment increased foci presence (mean ~8, P<0.0001 compared to NT, Figure 7). Neither PARPi nor PARGi treatment significantly increased the number of RAD51 foci (mean ~2 *vs.* ~4 *vs.* ~3, for PARPi veliparib, PARPi olaparib, and PARGi, respectively).

Altogether these data show that in response to PARPi, the HR DNA repair pathway, but not the NHEJ DNA repair pathway, is activated in PTEN-wildtype cells.

### PARPi and PARGi treatment do not induce FANCD2 foci accumulation in GBM cells

PARylation plays a role in the response of replication stress by participating in the repair of DNA damage, including both single-strand breaks and double-strand breaks and the resolution of stalled replication forks [16, 28]. Because the G2/M arrest observed in response to PARPi and PARGi could have resulted from unreplicated DNA, FANCD2, a marker of stalled replication forks, was analyzed (Figure 8, [29]). Foci were too small and often too numerous to count; therefore, cells were categorized as “positive” when FANCD2 foci were observed and “negative” when no foci were detected.

**Figure 8:**
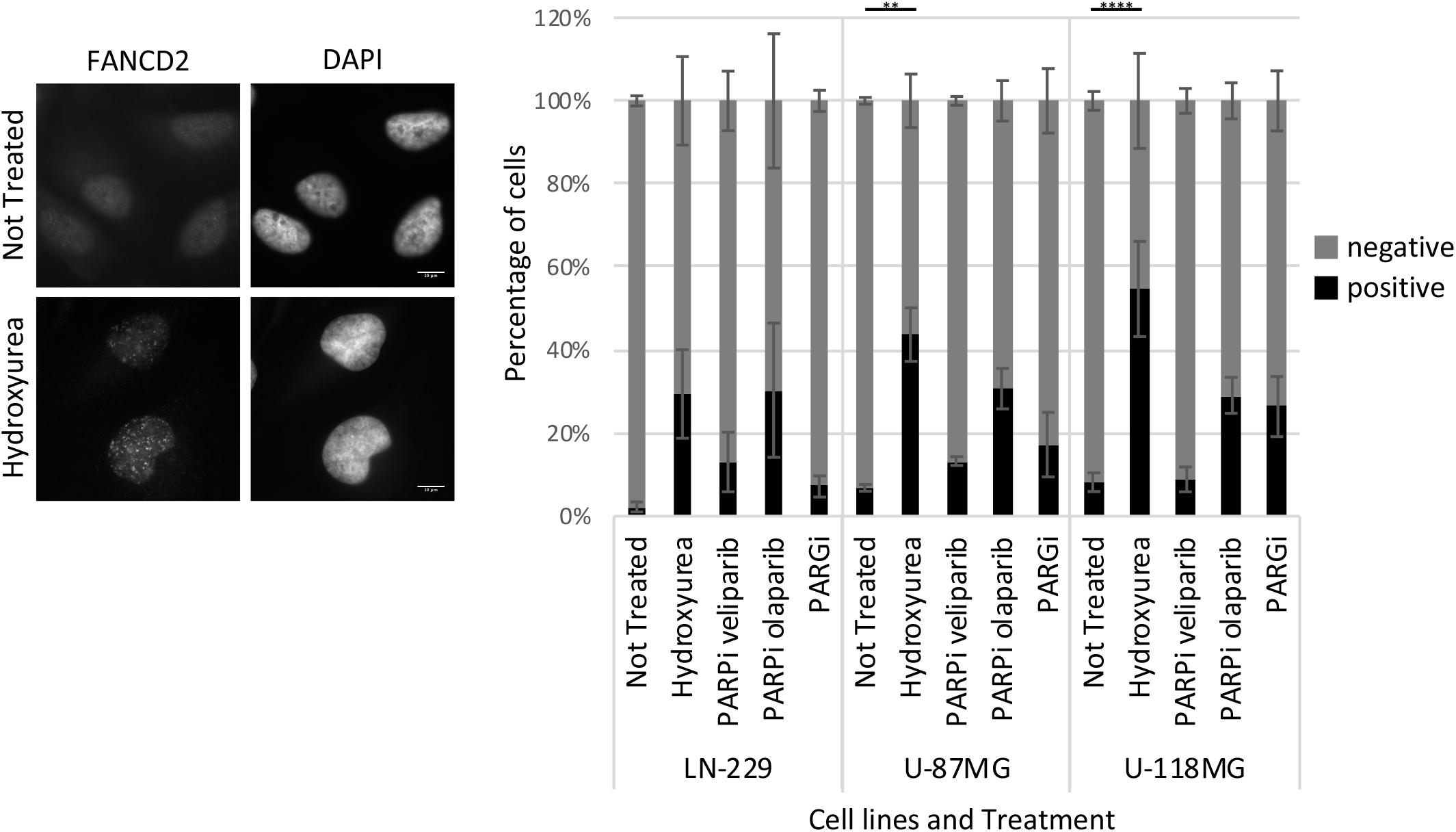
PARPi and PARGi are not associated with replicative stress in GBM cells. After 24 hours of treatment with either PARPi (veliparib or olaparib) or PARGi PDD00017273 at their IC_50_, GBM cells were fixed and immunofluorescence-stained for FANCD2. Nuclei were stained with DAPI. As a positive control, cells were treated with hydroxyurea. Representative images were presented (left panel). FANCD2 foci were examined in at least 100 cells and categorized as negative (black) or positive (medium gray). The average percentage of cells in each category was plotted for each cell line and treatment. Error bars represent standard deviation. Statistical significance was assessed by ANOVA. Data are representative of three biological replicates. **** P<0.0001; ** P≥0.0021.

**Figure 9:**
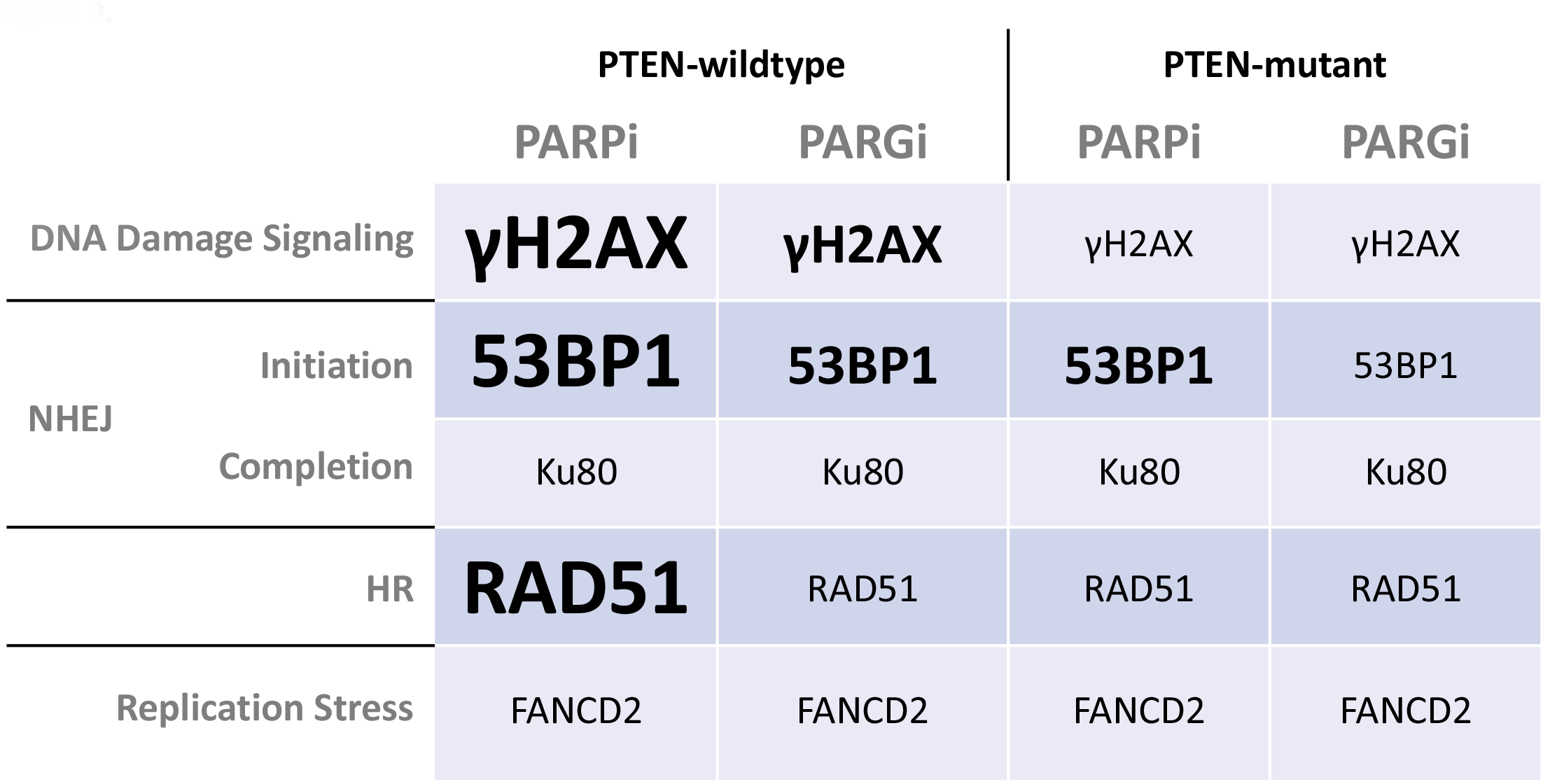
PARPi and PARGi differently affect the DNA damage response in PTEN-wildtype and PTEN-mutant GBM cell lines. γH2AX, 53BP1, Ku80, RAD51, and FANCD2 signals were assessed and summarized by PTEN status, and inhibitor treatment (PARPi *vs.* PARGi). Text size indicates foci formation response as compared to NT (small: treatment=NT; medium: treatment>NT; large: treatment≫NT).

Few LN-229 NT cells displayed FANCD2 foci (~2% of positive cells, Figure 8). In contrast, hydroxyurea treatment, which triggers replication stress, increased positive FANCD2 cells to ~30%. Neither PARPi (veliparib or olaparib) nor PARGi lead to the formation of FANCD2 foci.

FANCD2 foci were present in ~7% of U-87MG NT cells, and hydroxyurea treatment increased that percentage to ~45% (P=0.0011, Figure 8). Similar to LN-229 GBM cells, no statistically significant differences were measured between NT and PARPi (~15% positive cells for veliparib and ~30% positive cells for olaparib) or PARGi (~18% positive cells) treatment groups.

The same regulation observed in LN-229 and U-87MG cells was observed in U-118MG cells, with a basal level of ~9% of cells displaying FANCD2 foci and ~55% of cells having FANCD2 foci in response to hydroxyurea (P<0.0001, Figure 8). Similarly, neither PARPi (~9% positive cells for veliparib and ~30% positive cells for olaparib) nor PARGi (~30% positive cells) significantly changed FANCD2 foci number compared to non-treated cells.

These data suggest that neither PARP nor PARG inhibition induces replication stress in these GBM cells, regardless of PTEN status. Thus, the DNA damage (Figure 5) and the G2/M arrest previously observed (Figure 3) following treatment with these drugs do not likely result from DNA replication stress.

## Discussion

Here we report that GBM cells are differentially sensitive to PARP and PARG inhibition as a function of their PTEN status. However, no cell death is induced. Thus, these inhibitors in combination with PTEN loss may not be sufficient to induce synthetic lethality. This novel finding arose from the initial observation that PTEN-wildtype GBM cells were more sensitive to PARP inhibition, and PTEN-mutant cells to PARG inhibition.

The G2 arrest induced by PARP and PARG inhibition in PTEN-wildtype GBM cells (LN-229) and PTEN mutant GBM cells (U-87MG and U-118MG) may have resulted from the accumulation of DNA damage, preventing cells from completing aberrant chromosome segregation during mitosis. Analysis of the DNA damage biomarker γH2AX revealed an overall increase of DNA lesions upon PARPi, but not PARGi, in LN-229 GBM cells (wildtype PTEN). This result refutes our initial hypothesis and the general assumption that PTEN mutations may be associated with a HR-defective phenotype that lead to accumulation of DNA damage upon PARP inhibition, as observed in BRCA1/2-mutated breast and ovarian cancers [15]. The accumulation of γH2AX foci in LN-229 GBM cells treated with PARPi may be explained by either the direct generation of DNA damage by PARPi or defective DSB repair pathways.

NHEJ and HR are the major DNA DSB repair pathways. Their mechanism accounts for their dependence to cell cycle. PARP1 promotes both HR and alt-NHEJ, but inhibits classic NHEJ [19]. Thus, PARP inhibition would be expected to prevent the activation of HR and alt-NHEJ, and stimulate NHEJ. 53BP1 foci formation was analyzed to determine the repair pathway choice made by GBM cells in response to PARPi and PARGi. 53BP1 has no known enzymatic activity, yet by restricting end resection of DNA double-strand breaks in the G1 phase it promotes classic NHEJ. 53BP1 foci accumulation was associated with PARPi treatment in LN-229, but not in U-87MG or U-118MG cells, indicating that a PTEN mutation is not inevitably associated with a HR-defective phenotype and NHEJ activation. However, our data on the classic NHEJ mediator Ku80 suggest that final steps of this repair pathway are not-functional upon PARPi or PARGi in LN-229 cells.

Our data open the question on how DNA damages induced by PARPi are repaired. In PTEN wildtype GBM cells, the number of RAD51 foci was increased upon PARPi treatment, suggesting that HR is functional and activated. In contrast, RAD51 foci levels increased after PARPi olaparib in U-87MG. These observations suggest that the localization of PTEN mutations could modulate HR efficiency. The characterization of PTEN mutations that affect DSB repair by HR and contribute to BRCAness phenotype is essential for future research on GBM. While PARG inhibition mechanisms are currently unknown, its small effect on PTEN-mutant cells suggests that it is unlikely to function by molecular trapping. It also suggests that PARG inhibition is less sensitive to PTEN mutation differences. These data support the previously proposed hypothesis that PARPi and PARGi do not utilize the same mechanism.

The observation of G2 arrest without evidence for a cell death response may also result from replication stress in S phase. Thus, FANCD2 was assessed to investigate replication fork stalling. No significant changes in FANCD2 foci formation were observed following inhibitor treatments, indicating that the DNA damage accumulation and cycle arrest is not due to errors in DNA replication. The observed DNA damage is more likely the direct result of inhibitor effects.

Altogether, these results support that PARPi induce DNA damage in GBM cells that may be repaired by HR. Inhibitor efficacy has been found to vary in PTEN-mutant GBM, likely in function of mutation localization. In an attempt to personalize GBM treatment, GBM displaying mutations in the N-terminus of PTEN (U-87MG) may benefit from PARPi veliparib, while GBM with C-terminal mutations (U-118MG) may profit from both PARPi and PARGi treatment. Testing this new hypothesis in combination with current treatment will further characterize the molecular mechanism of SL in GBM for the long-term interest of patient survival.

## ACKNOWLEDGEMENTS

We thank members of our lab for technical help, discussion and critical feedback, including Trevor Wolf and Ross Cullen. We especially thank Dr. Diane Jaworski for sharing cell lines, and discussions on glioblastoma biology and Dr. Alissa Thomas for her clinical expertise on glioblastoma therapies.

Flow cytometry was performed at the Harry Hood Bassett Flow Cytometry and Cell Sorting Facility at the University of Vermont Larner College of Medicine for the use of BD LSRII. We would also like to thank Dr. Roxana del Rio-Guerra for assistance with the flow cytometry experiments.

Gene and protein expression data were acquired at the COBRE Neuroscience Cellular and Molecular Core at the University of Vermont Larner College of Medicine. This core is supported by the National Institute of General Medical Sciences of the National Institutes of Health (award number P30 RR 032135 / P30 GM 103498).

## Supplemental Tables

**Supplemental Table 1:** Distribution of cells (average percentage and standard deviation) over the cell cycle in function of treatment. TMZ: Temozolomide.

|  |  | Sub-G1 |  | G1 |  | S |  | G2 |  | Polyploidy |  |
| --- | --- | --- | --- | --- | --- | --- | --- | --- | --- | --- | --- |
|  |  | Average | Std dev | Average | Std dev | Average | Std dev | Average | Std dev | Average | Std dev |
| LN-229 | Not Treated | 2.5 | 0.2 | 55.9 | 0.1 | 22.9 | 0.0 | 14.5 | 0.5 | 4.4 | 0.7 |
|  | Digitonin | 1.5 | 0.1 | 57.6 | 1.7 | 23.5 | 0.9 | 13.7 | 0.9 | 3.8 | 0.2 |
|  | Doxorubicin | 1.9 | 0.2 | 58.8 | 0.6 | 23.0 | 0.3 | 13.3 | 0.3 | 3.1 | 0.4 |
|  | TMZ | 1.8 | 0.2 | 57.0 | 1.3 | 21.7 | 0.5 | 14.9 | 1.1 | 4.8 | 0.0 |
|  | PARPi veliparib | 2.7 | 0.1 | 26.9 | 0.5 | 12.4 | 1.2 | 51.3 | 1.8 | 6.8 | 0.1 |
|  | PARPi olaparib | 7.6 | 4.4 | 22.9 | 1.6 | 14.6 | 0.0 | 47.0 | 4.4 | 8.0 | 1.5 |
|  | PARGi | 2.6 | 0.2 | 64.9 | 1.1 | 13.4 | 2.7 | 15.2 | 2.0 | 4.0 | 0.6 |
| U-87MG | Not Treated | 2.5 | 0.3 | 25.8 | 0.4 | 5.6 | 0.3 | 36.7 | 0.4 | 29.6 | 0.8 |
|  | Digitonin | 3.3 | 0.0 | 26.3 | 0.0 | 6.7 | 0.0 | 38.3 | 0.0 | 25.4 | 0.0 |
|  | Doxorubicin | 2.5 | 0.5 | 26.7 | 0.9 | 8.4 | 0.9 | 36.3 | 0.4 | 26.2 | 0.2 |
|  | TMZ | 2.6 | 0.3 | 25.5 | 0.3 | 5.8 | 0.0 | 38.6 | 0.1 | 27.6 | 0.6 |
|  | PARPi veliparib | 5.6 | 0.0 | 16.4 | 1.2 | 3.7 | 0.2 | 36.4 | 0.3 | 38.1 | 0.7 |
|  | PARPi olaparib | 2.4 | 0.1 | 30.6 | 0.8 | 5.2 | 0.0 | 40.3 | 0.1 | 21.6 | 0.9 |
|  | PARGi | 2.5 | 0.1 | 28.1 | 1.4 | 4.4 | 0.3 | 47.0 | 1.5 | 18.2 | 0.2 |
| U-118MG | Not Treated | 3.4 | 0.6 | 39.4 | 2.2 | 10.5 | 0.4 | 31.3 | 0.3 | 15.6 | 1.0 |
|  | Digitonin | 32.6 | 0.3 | 29.5 | 0.3 | 8.4 | 0.2 | 22.7 | 0.1 | 6.9 | 0.3 |
|  | Doxorubicin | 4.2 | 0.8 | 38.7 | 1.1 | 9.9 | 0.6 | 32.2 | 0.1 | 15.3 | 1.1 |
|  | TMZ | 2.8 | 0.6 | 40.6 | 1.6 | 9.2 | 0.7 | 30.5 | 0.3 | 17.0 | 1.5 |
|  | PARPi veliparib | 3.0 | 0.8 | 43.7 | 0.9 | 9.3 | 0.8 | 29.8 | 1.9 | 14.3 | 1.1 |
|  | PARPi olaparib | 3.9 | 1.1 | 39.8 | 0.6 | 8.9 | 0.0 | 32.5 | 0.6 | 15.1 | 1.1 |
|  | PARGi | 4.0 | 0.5 | 30.1 | 0.9 | 8.9 | 1.1 | 41.3 | 0.3 | 15.9 | 0.5 |

**Supplemental Table 2:** Analysis of RAD51 foci.

|  | **LN-229** |  |  |  |  |
| --- | --- | --- | --- | --- | --- |
| **Nb Foci** | **Not-Treated** | **Etoposide** | **PARPi Veliparib** | **PARPi Olaparib** | **PARGi** |
| **min** | 0 | 0 | 0 | 0 | 0 |
| **max** | 56 | 99 | 110 | 124 | 32 |
| **median** | 0 | 1 | 1 | 4 | 0 |
| **mean** | 2 | 13 | 13 | 14 | 2 |

|  | **U-87MG** |  |  |  |  |
| --- | --- | --- | --- | --- | --- |
| **Nb Foci** | **Not-Treated** | **Etoposide** | **PARPi Veliparib** | **PARPi Olaparib** | **PARGi** |
| **min** | 0 | 0 | 0 | 0 | 0 |
| **max** | 34 | 102 | 76 | 125 | 70 |
| **median** | 0 | 0 | 0 | 1 | 0 |
| **mean** | 2 | 14 | 3 | 16 | 6 |

|  | **U-118MG** |  |  |  |  |
| --- | --- | --- | --- | --- | --- |
| **Nb Foci** | **Not-Treated** | **Etoposide** | **PARPi Veliparib** | **PARPi Olaparib** | **PARGi** |
| **min** | 0 | 0 | 0 | 0 | 0 |
| **max** | 30 | 56 | 53 | 68 | 41 |
| **median** | 0 | 0 | 0 | 0 | 0 |
| **mean** | 1 | 8 | 2 | 4 | 3 |

